# Nerve injury inhibits *Oprd1* and *Cnr1* transcription through REST in primary sensory neurons

**DOI:** 10.1101/2024.02.17.579842

**Authors:** Ashok Subedi, Aadhya Tiwari, Asieh Etemad, Yuying Huang, Biji Chatterjee, Samantha M. McLeod, Yungang Lu, DiAngelo Gonzalez, Krishna Ghosh, Sanjay K. Singh, Elisa Ruiz, Sandra L. Grimm, Cristian Coarfa, Hui-Lin Pan, Sadhan Majumder

## Abstract

The transcription repressor REST in the dorsal root ganglion (DRG) is upregulated by peripheral nerve injury and promotes the development of chronic pain. However, the genes targeted by REST in neuropathic pain development remain unclear. The expression levels of 4 opioid receptor (*Oprm1*, *Oprd1*, *Oprl1*, *Oprk1)* and the cannabinoid CB1 receptor (*Cnr1*) genes in the DRG regulate nociception. In this study, we determined the role of REST in the control of their expression in the DRG induced by spared nerve injury (SNI) in both male and female mice. Transcriptomic analyses of male mouse DRGs followed by quantitative reverse transcription polymerase chain reaction analyses of both male and female mouse DRGs showed that SNI upregulated expression of *Rest* and downregulated mRNA levels of all 4 opioid receptor and *Cnr1* genes, but *Oprm1* was upregulated in female mice. Analysis of publicly available bioinformatic data suggested that REST binds to the promoter regions of *Oprm1* and *Cnr1*. Chromatin immunoprecipitation analyses indicated differing levels of REST at these promoters in male and female mice. Full-length *Rest* conditional knockout in primary sensory neurons reduced SNI-induced pain hypersensitivity and rescued the SNI-induced reduction in the expression of *Oprd1* and *Cnr1* in the DRG in both male and female mice. Our results suggest that nerve injury represses the transcription of *Oprd1* and *Cnr1* via REST in primary sensory neurons and that REST is a potential therapeutic target for neuropathic pain.

## Introduction

Chronic neuropathic pain is difficult to treat and remains a major clinical problem, adversely affecting the quality of life of millions of Americans (Finnerup et al. 2016; Fullerton, Doyle, and Murphy 2018; Fricker et al., 2020; Ghosh and Pan 2022; Thouaye and Yalcin 2023). The mechanisms underlying the transition from acute to chronic pain after nerve injury remain unclear, but mounting evidence suggests that epigenomic mechanisms (an array of modifications, including DNA methylation and histone marks) play a major role in this transition (Descalzi et al. 2015; Ghosh and Pan 2022). Persistent changes in the upregulation of pro-nociceptive genes and downregulation of anti-nociceptive genes account for the long-lasting aberrant excitability of dorsal root ganglion (DRG) neurons and excitatory nociceptive input to the spinal cord (Wang et al. 2002; Laumet et al. 2015; Garriga et al. 2018). For example, nerve injury causes upregulation of α2δ-1 in the DRG, which leads to increased synaptic NMDA receptor activity and glutamatergic input to spinal dorsal horn neurons (Chen et al. 2018; Zhang et al. 2022). Furthermore, nerve injury reduces the expression of almost all voltage-gated K^+^ channels (Rasband et al. 2001; Chen, Cai, and Pan 2009b; Rose et al. 2011; Laumet et al. 2015), contributing to the increased excitability of DRG neurons. Histone methyltransferase G9a-mediated epigenetic regulation in DRG neurons is also essential for the acute-to-chronic pain transition after nerve injury (Laumet et al. 2015).

Opioids, the most powerful analgesics, act through opioid receptors, which belong to the G-protein coupled receptor family (MOR for μ, KOR for κ, DOR for δ, NOP for nociceptin/orphan FQ or N/OFQ) (Fricker et al., 2020). However, chronic opioid treatment causes analgesic tolerance/hyperalgesia, which has resulted in an opioid addiction epidemic in the U.S. The various opioid receptors have been shown to have diverse roles in opioid sensitivity and the development of opioid tolerance. In this regard, opioid-induced analgesia and hyperalgesia/analgesic tolerance depend critically on the expression level of O*prm1*, which encodes the μ-opioid receptor, in DRG neurons (Sun et al. 2019). Also, the δ-opioid receptor (DOR) in primary sensory neurons regulates the analgesic effect of DOR agonists but not morphine-induced analgesic tolerance (Jin et al. 2022). Nerve injury in male mice reduces opioid analgesia via G9a-mediated repression of *Oprm1* (Zhang et al. 2016). G9a also suppresses cannabinoid CB1 receptor (encoded by *Cnr1*) expression in primary sensory neurons (Luo et al. 2020). In addition, peripheral nerve injury causes overexpression of the transcriptional repressor RE1-silencing transcription factor (REST, also known as Neuron Restrictive Silencer Factor) (Kagalwala, Singh, and Majumder 2008; Gopalakrishnan 2009; Nechiporuk et al. 2016) in the DRG of male mice, which represses REST target genes such as *Kcnd3, Kcnq2, Scn10a, Chrm2,* and the *Oprm1* (Willis et al. 2016; Zhang et al. 2018). However, it remains unclear whether REST controls the expression of the 4 opioid receptor and CB1 receptor genes in the DRG.

All previous studies of REST-mediated chronic pain (Zhang et al. 2018; Zhang et al. 2019) were performed using a *Rest* conditional knockout (cKO) mouse model that has now been discovered to harbor only partial *Rest* cKO (Nechiporuk et al. 2016) with only ∼40% loss of REST activity, generating confusion in some areas of research on primary sensory neurons. The Mandel lab then created a highly robust full-length *Rest* cKO mouse line (Nechiporuk et al. 2016), which was used in the present study. To our knowledge, this would be the first use of the full-length *Rest* cKO mouse line in a chronic pain study. In addition, most of the previous studies using the *Rest* cKO mouse model were performed with male mice only. Thus, here we studied the role of REST in nerve injury-induced chronic pain (NICP) in controlling the expression of opioid and cannabinoid CB1 receptor genes in DRG neurons in both male and female mice.

## Results

### Peripheral nerve injury induces chronic pain hypersensitivity similarly in male and female wild-type mice

Most previous studies using mouse models of neuropathic pain used only male mice. To determine whether there is a sex difference in neuropathic pain development, we performed either spared nerve injury (SNI) or sham surgery in both male and female wild-type C57BL/6 mice. We then measured ipsilateral hindpaw withdrawal thresholds in response to a noxious pressure stimulus (mechanical hyperalgesia), noxious radiant heat (thermal hyperalgesia), and von Frey filaments (tactile allodynia) for 4 weeks (Figure 1). SNI induced similar reductions in the withdrawal thresholds in the injured hindpaws of both male and female mice. Mechanical and thermal hypersensitivity developed within 5 days and persisted for at least 4 weeks in mice of both sexes.

**Figure 1.**
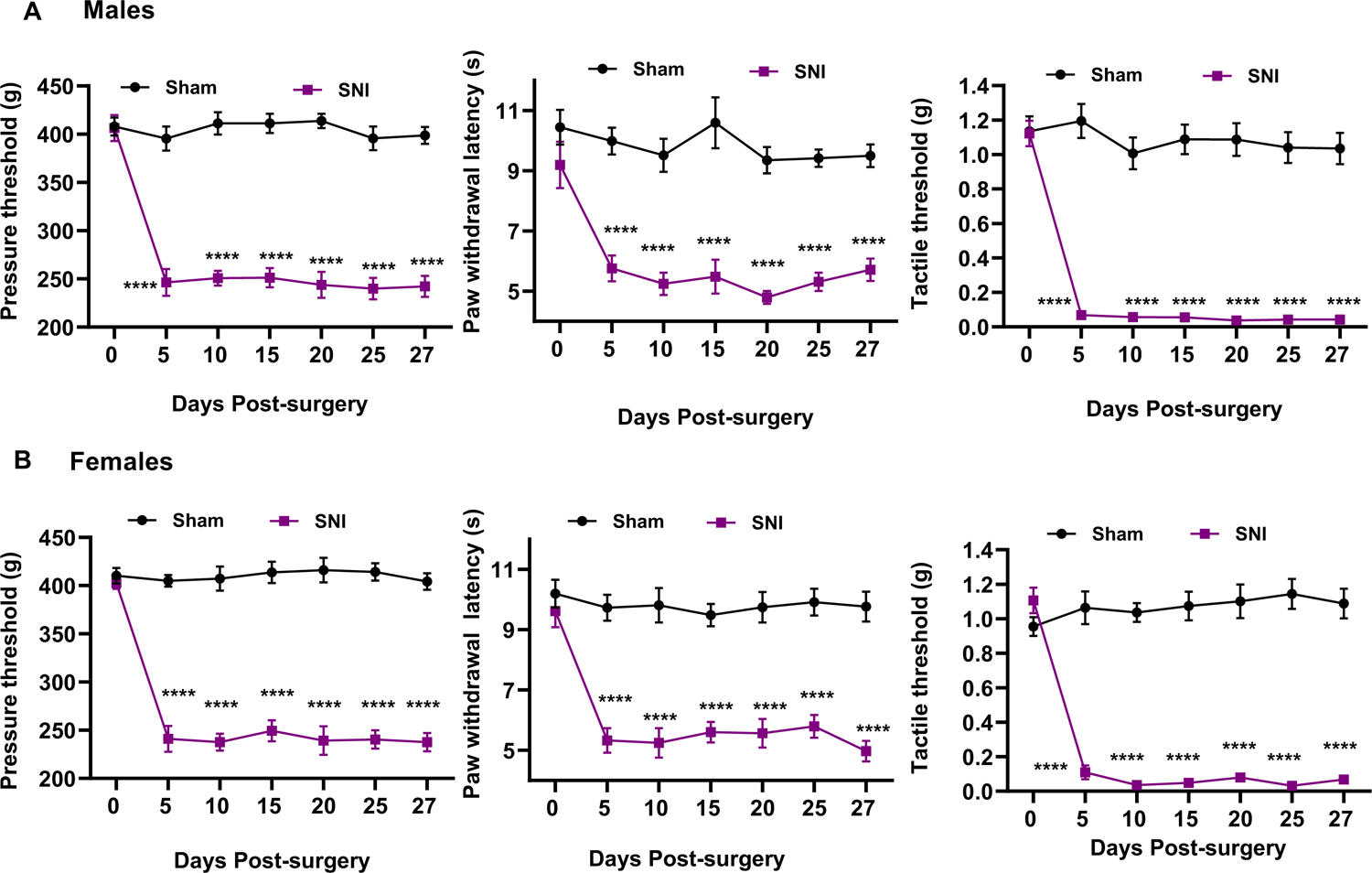
Peripheral nerve injury induces chronic neuropathic pain hypersensitivity similarly in male and female wild-type mice. Either spared nerve injury (SNI) or sham surgery (Sham) in the left hind limbs of both male and female wild-type mice was performed. Pain symptoms in terms of ipsilateral hindpaw withdrawal thresholds (withdrawal latency) were determined in response to a noxious pressure stimulus (mechanical hyperalgesia), noxious radiant heat (thermal hyperalgesia), and von Frey filaments (tactile allodynia) for 29 days. n=6 per group. 2 way ANOVA test was used to compare groups. Data are presented as mean+-SEM.

Because we measured evoked pain hypersensitivity using a reflex response, we performed rotarod tests to determine whether SNI induces the impairment of motor functions. As shown in Supplementary Figure S1, there were no significant differences in the rotarod performance tests in mice treated with SNI or sham surgery in either sex.

### Nerve injury leads to transcriptomic activation of *Rest* and repression of opioid receptor and cannabinoid CB1 receptor genes in the injured DRG of wild-type mice

Previous studies showed that nerve injury impacts expression of *Oprm1, Oprd1, and Cnr1* in the DRG (Luo et al. 2002; Sun et al. 2019; Luo et al. 2020). To determine whether other opioid receptors are also impacted by nerve injury, we collected ipsilateral DRG tissues at the L3/L4 level 28 days after SNI or sham surgery, as shown in Figure 1, from individual mouse. We then extracted mRNA from the tissues, picked 3 individual mRNA samples at random, and performed RNA sequencing. We analyzed the data as we described previously (Marisetty et al. 2017; Lu et al. 2018; Marisetty et al. 2019).

As shown in Figure 2, 1261 genes were differentially expressed between the sham-surgery and SNI groups. In terms of opioid receptors and cannabinoid receptors, a robust separation between sham-surgery and SNI groups was observed (Figure 2A). Volcano plots showed significant upregulation of 1058 genes and downregulation of 203 genes in the injured DRG (Figure 2B). While the *Rest* gene transcript was induced by SNI, opioid receptor genes *Oprd1*, *Oprm1*, *Oprl1*, and *Oprk1,* as well as cannabinoid CB1 receptor gene *Cnr1,* were suppressed (Figure 2C). GSEA corroborated this observation, showing suppression of opioid signaling and G-protein coupled opioid receptor signaling pathways (Figure 2D). These results suggest that nerve injury represses the expression of all 4 opioid receptor and *Cnr1* genes in the DRG of male mice.

**Figure 2.**
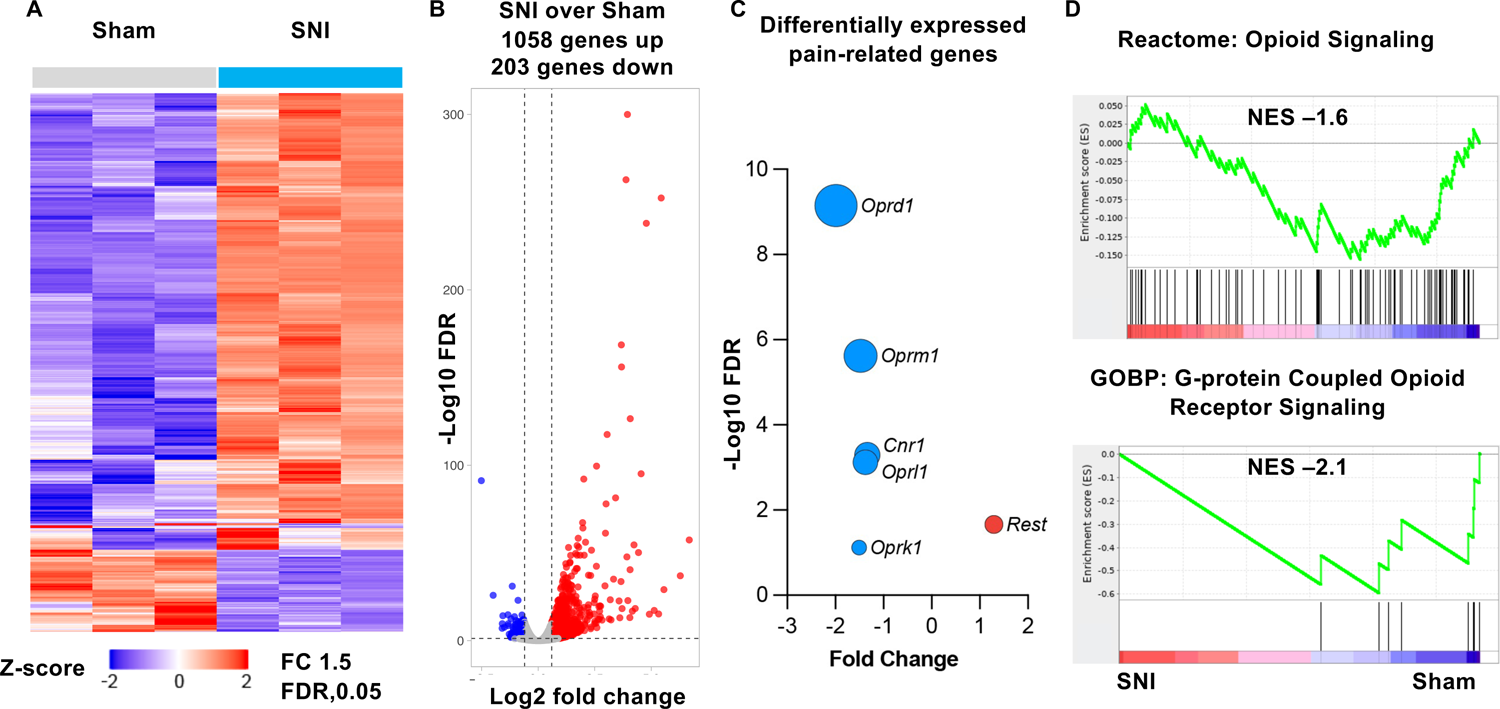
SNI leads to transcriptomic activation of *Rest* and repression of opioid receptor and cannabinoid CB1 receptor genes in the injured DRG of male wild-type mice. A. Hierarchical clustering of differentially expressed genes between SNI and Sham surgery groups. B. Volcano plot showing differentially expressed genes opioid receptor and cannabinoid receptor genes. C. *Rest*, opioid receptor genes, and cannabinoid receptor gene expression in SNI and Sham surgery groups. D. Gene Set Enrichment Analysis shows downregulation of the opioid signaling pathway. Experiments were performed in triplicate; L3 and L4 DRGs from 3 individual mice were examined.

### SNI increases mRNA levels of *Rest* and decreases mRNA levels of opioid receptor and cannabinoid CB1 receptor genes in the DRG of both male and female wild-type mice

We then used RT-qPCR to analyze the mRNA levels of *Rest* and the 4 opioid receptor and *Cnr1* genes in L3/L4 DRG samples from male mice 28 days after SNI surgery. As shown in Figure 3A, nerve injury significantly increased the expression of *Rest* as previously shown by others (Zhang et al. 2018; Zhang et al. 2019). Indeed, Western blotting assays indicated a corresponding increase in the REST protein (Supplementary Figure S2). Further, nerve injury significantly decreased the expression of *Oprm1*, *Oprd1*, *Oprk1*, *Oprl1*, and *Cnr1* transcripts (Figure 3A), validating the RNA-Seq results.

**Figure 3.**
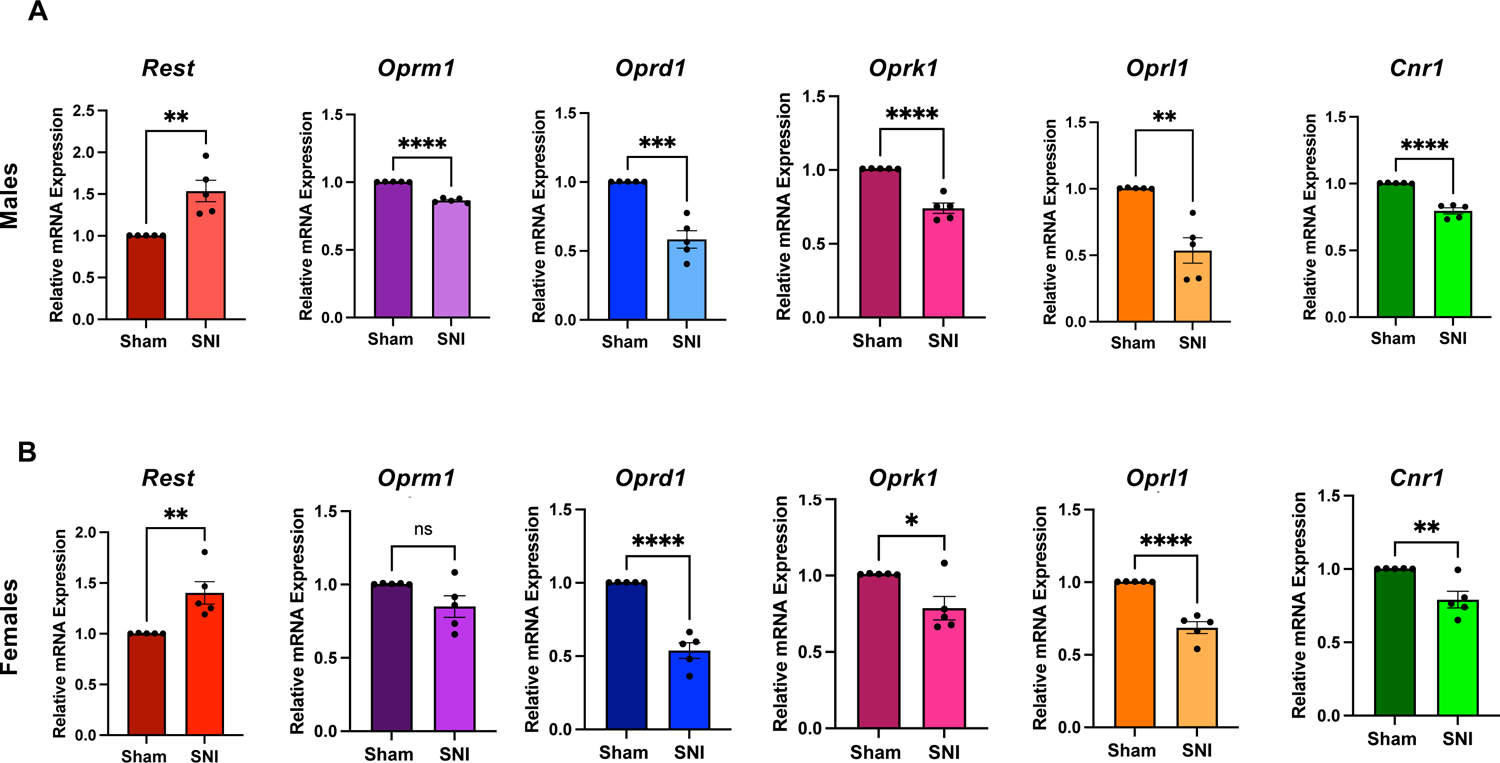
SNI results in increased mRNA levels of *Rest* and decreased mRNA levels of opioid receptor and cannabinoid CB1 receptor genes in the injured DRG of both male and female wild-type mice. mRNA samples from the L3/L4 DRG tissues of (A) male and (B) female mice characterized in Figure 1, were used for RT-qPCR analysis of *Rest*, *Oprm1*, *Oprd1*, *Oprk1*, *Oprl1*, and *Cnr1* transcripts. Student t test was used to measure the p-value. n=5 mice per group. Data are presented as mean+-SEM.

To determine whether SNI has a similar effect on the expression of these receptor genes in DRG in female mice, we performed similar experiments in female mice. As shown in Figure 3B, SNI increased *Rest* and decreased *Oprd1*, *Oprk1*, *Oprl1*, and *Cnr1* transcripts in the DRG of female mice. Unexpectedly, SNI did not show any significance difference in the *Oprm1* transcript levels in the DRG of female mice.

### REST binds differently at the *Oprm1* and *Cnr1* promoters in the injured DRG in male and female wild-type mice

We used publicly available bioinformatic data using the geneXplain platform (https://genexplain.com/genexplain-platform/) to identify potential REST binding sites on the promoter regions of *Oprm1* and *Cnr1*. To determine whether nerve injury affects the REST binding at the *Oprm1* and *Cnr1* promoters, we performed chromatin immunoprecipitation (ChIP)-qPCR analysis of L3/L4 DRG samples from male and female mice 28 days after SNI (Figure 4). SNI increased the presence of REST on the *Oprm1* promoter in the DRG of male mice but did not show any significant difference in female mice. The ChIP data corroborated our RT-qPCR results, suggesting that REST in the DRG plays a role in repressing *Oprm1* transcript levels in male but not in female mice after nerve injury.

**Figure 4.**
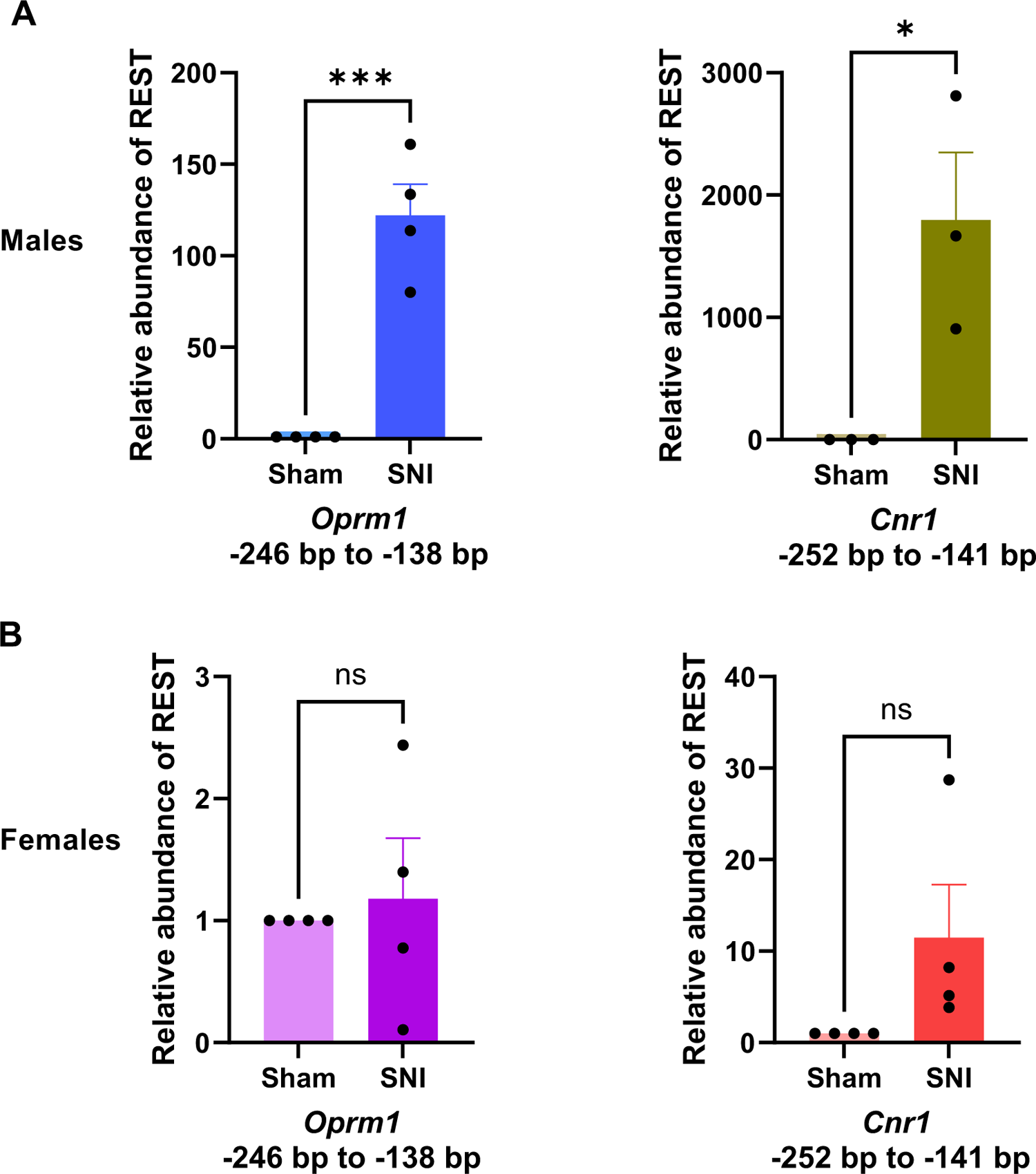
REST binds differently at the *Oprm1* and *Cnr1* promoters in the injured DRG in male and female wild-type mice. REST binding to *Oprm1* and *Cnr1* promoters was measured by qChip assay in sham and SNI in (A) male and (B) female mice. n= 5 animals per group. Student t test was used to measure the p-value. Data is presented as Mean+-SEM

Furthermore, ChIP-qPCR analysis showed that SNI increased REST binding at the *Cnr1* promoter in the DRG of both male and female mice. However, whereas this binding was statistically significant in the male mice, it did not reach significance in the female mice (Figure 4).

### DRG neuron-specific *Rest* cKO attenuates nerve injury-induced pain hypersensitivity in both male and female mice

Next, we used full-length *Rest* cKO mice (Nechiporuk et al. 2016) to determine whether the increased expression of *Rest* in DRG neurons has a causal role in the development of chronic neuropathic pain. We performed SNI or sham surgery in male *Rest* cKO mice, which included 3 groups: Control (-Cre) + sham, control (-Cre) + SNI, and *Rest* cKO (+Cre) + SNI. We measured pressure, heat, and tactile withdrawal thresholds in the ipsilateral hindpaws for 4 weeks, as we did in the wild-type mice. As shown in Figure 5A, SNI profoundly reduced pressure, thermal, and tactile withdrawal thresholds of the ipsilateral hindpaw in the control (-Cre) + SNI mice compared with the control (-Cre) + sham mice. The pressure and thermal withdrawal thresholds of the ipsilateral hindpaw did not differ significantly between the male *Rest* cKO and control littermate groups during the first 3 days after SNI (corresponding to the acute phase of pain). However, SNI-induced pressure and heat hypersensitivities were markedly attenuated in *Rest* cKO mice 6 to 26 days after SNI; tactile hypersensitivity was attenuated in *Rest* cKO mice 8 to 26 days after SNI (a period corresponding to the transition to the chronic phase of pain). These results were similar to those we observed using a different strain of male *Rest* cKO mice (Zhang et al. 2018).

**Figure 5.**
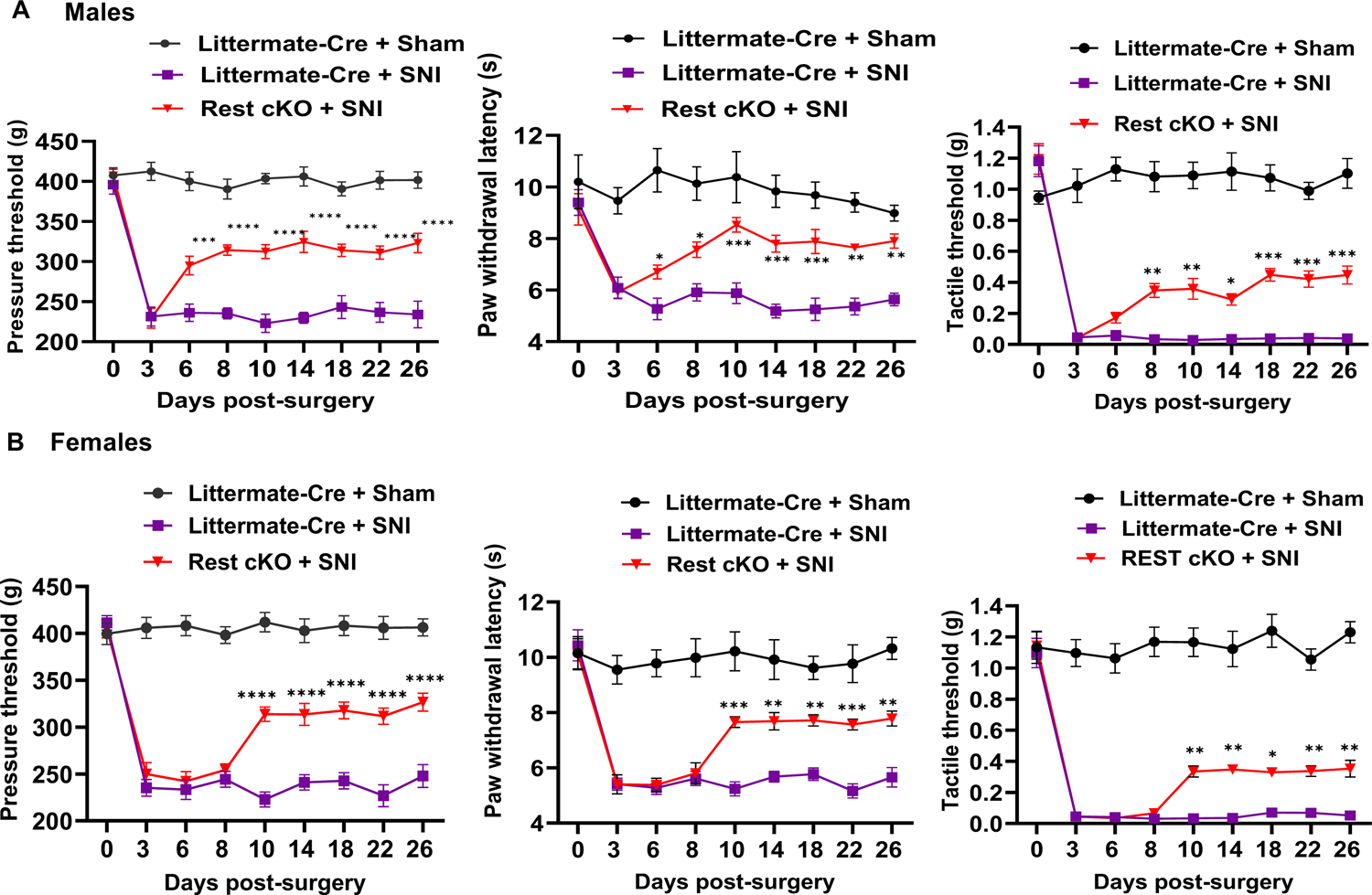
DRG neuron-specific *Rest* cKO attenuates nerve injury-induced pain hypersensitivity in both male and female mice. Nerve injury-induced pain hypersensitivity was assessed in (A) male and (B) female mice as described in Figure 1. The mouse groups were: littermate (-Cre) (control) + Sham; littermate (-Cre) (control) + SNI; and *Rest* cKO (+Cre) + SNI. n=6 per group; two-way ANOVA followed by Tukey’s multiple comparison test. Data are presented as mean+/-SEM.

We then performed similar experiments using female *Rest* cKO mice (Figure 5B). The female control (-Cre) mice, like the males, showed a profound reduction in pressure, thermal, and tactile withdrawal thresholds after SNI. All three withdrawal thresholds of the ipsilateral hindpaw did not differ significantly between the female *Rest* cKO (+Cre) and control littermate (-Cre) groups during the first 8 days after SNI. These hypersensitivities were then markedly mitigated in *Rest* cKO mice 10 to 26 days after SNI. These results suggest that REST in DRG neurons contributes to nerve injury-induced chronic pain in both male and female mice. Further, the slight delay in onset of pressure, thermal, and tactile hypersensitivities in female *Rest* cKO mice, as compared to the male cKO mice, suggests that REST may have a more significant role in promoting chronic pain development in male than in female mice.

We then performed rotarod tests to determine whether SNI induces the impairment of motor functions. As shown in Supplementary Figure S3, there were no significant differences in the rotarod performance tests in *Rest* cKO or littermate mice treated with SNI or sham surgery in either sex.

### REST represses the expression of *Oprd1* and *Cnr1* in DRG neurons of nerve-injured male and female mice

We then determined the role of *Rest* in DRG neurons in SNI-induced downregulation of opioid and cannabinoid receptor genes in male *Rest* cKO mice. We performed RT-qPCR using L3/L4 DRG tissues from control and *Rest* cKO mice 28 days after SNI or sham surgery. As shown in Figure 6A. SNI in male controls increased expression of *Rest* and decreased expression of *Oprm1*, *Oprd1*, *Oprk1*, *Oprl1,* and *Cnr1* like what we saw with wild-type mice in Figure 3. As expected, the *Rest* expression level in the DRG was much lower in *Rest* cKO + SNI mice than in control + SNI mice. However, we did not observe complete elimination of *Rest* in the DRG in *Rest* cKO mice. This observation is similar to the previous observations reported earlier using a different *Rest* cKO mouse line (Zhang et al. 2018; Zhang et al. 2019). One of the reasons for these results is likely because Advillin-Cre–induced target gene knockout does not occur in 100% of DRG neurons (Zappia et al. 2017). Remarkably, the diminished expression levels of *Oprd1* and *Cnr1* in control + SNI mice were significantly restored in *Rest* cKO + SNI mice. However, the expression levels of *Oprm1, Oprk1,* and *Oprl1* in the DRG did not differ significantly between *Rest* cKO + SNI and control + SNI mice.

**Figure 6.**
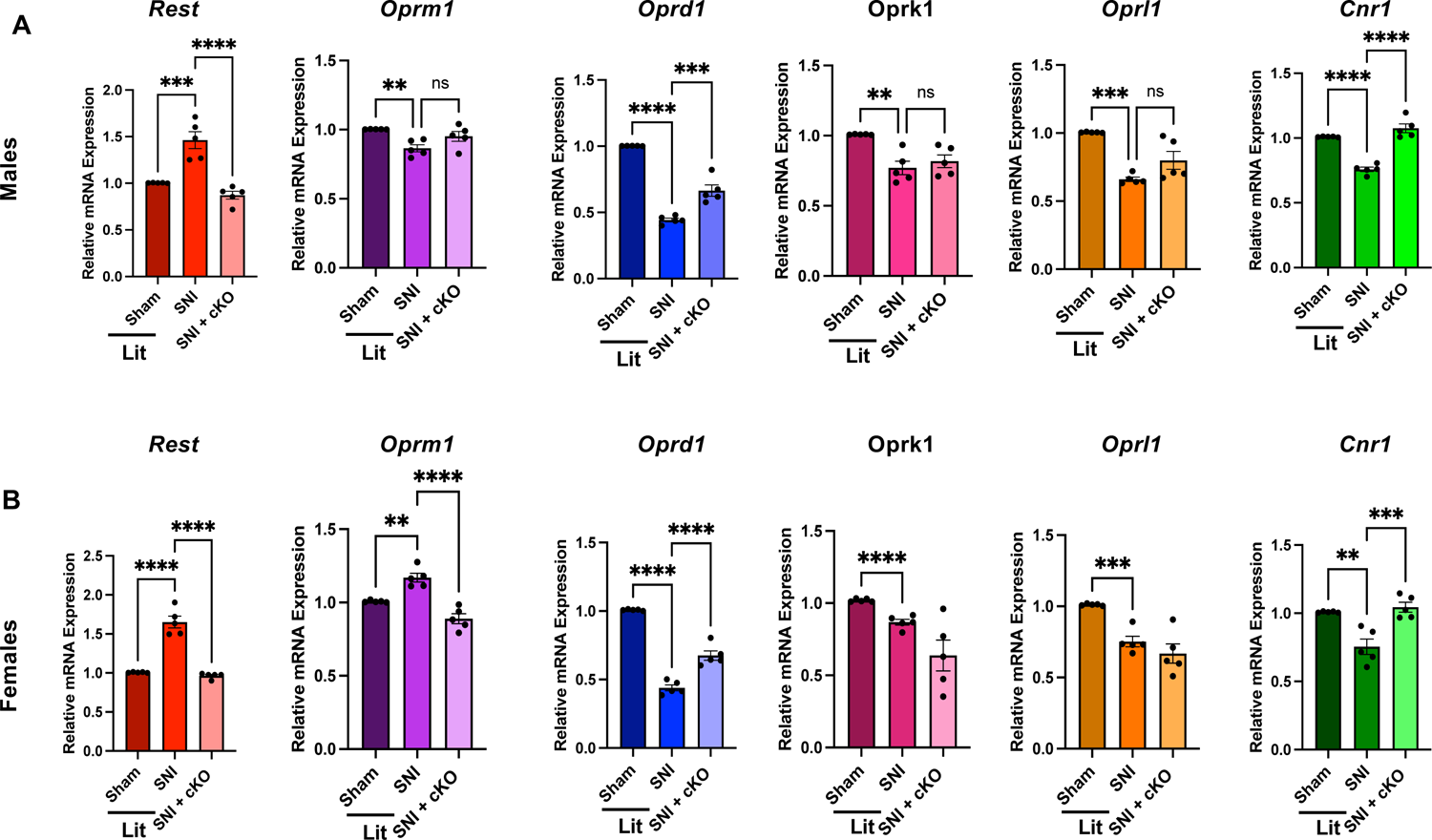
REST represses the expression of *Oprd1* and *Cnr1* in DRG neurons of nerve-injured male and female mice. mRNA samples from the L3/L4 DRG tissues of (A) male and (B) female mice characterized in Figure 5, were used for RT-qPCR analysis of *Rest*, *Oprm1*, *Oprd1*, *Oprk1*, *Oprl1*, and *Cnr1* transcripts. Lit=littermates. P-values are shown. n=5 mice per group. Data are presented as mean+-SEM.

In female control littermate mice, SNI also significantly increased the mRNA level of *Rest* in the DRG and reduced the mRNA levels of *Oprd1*, *Oprk1*, *Oprl1,* and *Cnr1* in the DRG (Figure 6B). In contrast to males, however, SNI increased the mRNA level of *Oprm1* in the DRG of female mice (Figure 6B). Like male mice, *Rest* cKO in females, as compared to control littermate + SNI, significantly reversed the reduced expression levels of *Oprd1* and *Cnr1* in the DRG. However, unlike its effect in males, SNI significantly increased the mRNA level of *Oprm1* in the DRG in the female littermate controls, and this increase was reversed in female *Rest* cKO + SNI mice. Taken together, our results suggest that *Oprd1* and *Cnr1* genes were robustly silenced by nerve injury via REST in primary sensory neurons of both male and female mice.

## Discussion

Our studies suggest that the impact of nerve injury to the DRG induced REST-mediated reduction of the *Oprm1* transcript levels, measured at day 28 after injury, of male but not female mice. These transcript levels were confirmed by our ChIP data showing the presence of REST on the *Oprm1* gene chromatin in males but not in females. One possible reason for this difference could be the timing of the assay. As previously reported (Lee et al. 2011), expression of the *Oprm1* gene in primary afferent neurons after spinal nerve injury is time dependent. A second possibility is that sex hormones affect REST binding to the *Oprm1* gene chromatin in female mice (Sharp, Pearson, and Smith 2022). This might suggest that morphine-induced analgesia would be more effective in the DRG of female mice than in male mice. Interestingly, one study found that, after surgery, women used lower dosage of opioids than men (Gear et al. 2003). However, several confounding factors could account for that observation. Indeed, reports of sex differences in opioid analgesia have been controversial (Craft 2003; Fillingim and Gear 2004; Niesters et al. 2010). In addition, the actual impact of MOR is likely regulated by MOR-DOR heteromer formation (Gomes et al., 2016).

Our studies also indicated that REST in DRG neurons has a more prominent role in promoting the transition to chronic neuropathic pain in males than in females. The small delay in onset of pressure, thermal, and tactile hypersensitivities in *Rest* cKO female mice suggests that REST may have a more significant role in promoting chronic pain development in males than in females.

*Cnr1* expression levels in DRG neurons are important for regulating pain hypersensitivity; *Cnr1* cKO in DRG neurons causes pain hypersensitivity in mice (Liu et al. 2021). Similarly, we observed that nerve injury diminishes *Cnr1* expression in the DRG via G9a (Luo et al. 2020), a cofactor of REST. We found that *Rest* cKO similarly attenuated thermal, pressure, and touch hypersensitivities in the DRG in both male and female nerve-injured mice. This decrease in *Cnr1* expression also corresponded to the presence of REST protein on the *Cnr1* gene chromatin in the injured DRG. These results suggest that the REST-G9a complex is involved in this process. Further, *Oprd1* cKO in DRG neurons increases pain hypersensitivity caused by nerve injury or tissue inflammation in mice, indicating that DOR expression levels in DRG neurons normally restrain pain hypersensitivity (Gaveriaux-Ruff et al. 2011; Jin et al. 2022). Because there is no potential RE1-binding site at the *Oprd1* promoter, *Rest* likely regulates DOR expression in DRG neurons indirectly (e.g., by altering the expression of a transcription factor that binds to the *Oprd1* promoter or via one or more microRNA targets). We are not aware of any DRG cKO studies on *Oprk1* and *Oprl1*. However, constitutive KO of *Oprk1* in mice enhances sensitivity to chemically induced acute visceral pain (Simonin et al. 1998). *Oprl1* KO mice do not display differences in nociceptive thresholds when compared with wild-type animals (Nishi 1997; Mamiya et al. 1998). These studies suggest that nerve injury-induced *Rest* upregulation likely promotes pain hypersensitivity mainly via downregulation of *Oprd1* and *Cnr1* in DRG neurons.

Because REST cKO in DRG neurons attenuates SNI-induced pain hypersensitivity and reduction in the expression of *Oprd1* and *Cnr1*, it is conceivable that inhibiting REST activity would reduce chronic neuropathic pain and augment opioid/cannabinoid analgesic actions by increasing the expression of *Oprd1* and *Cnr1* genes in DRG neurons. Thus, REST might present a valuable target for treating neuropathic pain.

## Methods

### Mice

All mouse experiments were approved by the Institutional Animal Care and Use Committee (IACUC) of The University of Texas MD Anderson Cancer Center. We used C57BL/6 wild-type mice obtained from the Research Animal Support Facility at The University of Texas MD Anderson Cancer Center.

Wild-type C57B6 mice were purchased from in-house MDAnderson facility (Experimental Radiation Oncology) and then bred in our mouse colony. *Rest* conditional full-length KO (obtained from, mice were obtained from Dr. Gail Mandel (Nechiporuk et al. 2016), and characterized by measuring transcript and protein levels of the *Rest* gene in littermates (-Cre) and cKO (+Cre) mice. Deletion of *Rest* in mouse DRG neurons was generated by crossing female *Rest*-loxP^+/+^ mice and male Advillin-Cre mice (purchased form The Jackson Laboratory, Bar Harbor, ME, #032536). Advillin-Cre^+/-^: *Rest*-loxP^+/-^ mice obtained from the first generation were crossed with female *Rest*-loxP^+/+^ mice to get Advillin-Cre^+/-^: *Rest*-loxP^+/+^ mice (*Rest*-cKO mice). Littermates without Cre expression (Advillin-Cre^-/-^: *Rest*-loxP^+/+^) were used as controls. Mice were earmarked at the time of weaning (3 weeks after birth), and tail biopsies were used for polymerase chain reaction genotyping.

The primers used for genotyping Advillin-Cre^+/-^ were: Avil/003: CCCTGTTCACTGTGAGTAGG; Avil/002: AGTATCTGGTAGGTGCTTCCAG; Cre/01:GCGATCCCTGAACATGTCCATC. The wild-type allele produces a 500-bp product, and the mutant allele produces a 180-bp product. For genotyping *Rest-GTi/ GTi*, the primers used were: GTA5: TGGATGTTGAGGTCCGTTGTG, GTB5: GCTACGGATCCCTTCTTCCC, and GTB1: AACGGCCCCCGACGTCCCTGG to produce a wild-type product of 480 bp and a mutant product of 600 bp. In addition, Western blotting assays indicated substantial loss of REST protein in the DRG of both male and female *Rest* cKO mice compared to those of their littermates (Supplementary Figure S4).

Both male and female adult mice (N = 5-6 per group; 8-10 weeks of age) were used for the final experiments. To improve the breeding performance of *Rest* cKO mice, DietGel Prenatal supplement (ClearH_2_O, Inc., Westbrook, ME, #69-503-02) was given as a feed additive to the breeding pairs (10 g per mouse twice weekly).

Spared nerve injury. SNI surgery was performed as we and others have described previously (Laedermann et al. 2014; Laumet et al. 2015). Mice were anesthetized with 2-3% isoflurane, and the sciatic nerve and its 3 terminal branches (the sural, common peroneal, and tibial nerves) of the left leg were exposed under a surgical microscope. The tibial and common peroneal nerves were tightly ligated with a 6-0 silk suture and sectioned distal to the ligation sites, leaving the sural nerve intact. Mice were monitored every day for 3 days post-surgery and post-operative cards were filled every day as approved by the IACUC protocol. Mice did not receive opioid analgesia after surgery because it could confound the results.

### Nociceptive behavior test

We used hindpaw withdrawal thresholds in response to a noxious pressure stimulus (mechanical hyperalgesia), a noxious radiant heat stimulus (thermal hyperalgesia), and von Frey filaments (tactile allodynia) to assess pain, as we have published previously (Zhang et al. 2018; Jin et al. 2022). To familiarize the mice with the testing conditions and environments, they were kept on the wired metal grid and thermal testing apparatus for 30 minutes each day for 3 days before testing began.

A Rodent Pincher analgesia meter (IITC Life Science) was used to measure the pressure thresholds in the mice. The hindpaw was put between the 2 arms of the pincher, and steady pressure was applied at increasing intensity until the mouse showed a pain response, as indicated by paw withdrawal and/or vocalization. The force in “*g*” that elicited the pain response was recorded. Each trial was repeated 3 times with an interval of 2 minutes in between, and the mean value was recorded.

For thermal sensitivity, the plantar test (Hargreaves method) with heated glass (IITC Life Science, Woodland Hills, CA) was used. The glass surface was maintained at 30 °C, and the active intensity of the radiant lamp was maintained at 30%. Mice were acclimatized for 30 minutes in the observation chamber before testing. The planar surface was heated by a mobile radiant heat source, and the time of paw withdrawal was recorded by an automatic time recorder associated with the apparatus. Each trial was repeated 3 times, and the mean time to withdrawal was calculated.

For the tactile sensitivity testing, the mice were habituated on the wired metal grid for 30 minutes immediately before the test. The von Frey filament (Stoelting, Wood Dale, IL) was applied to the planar surface of the hindpaw for 3-5 seconds. Brisk withdrawal or paw licking was considered a positive response. The filament force value was increased until a positive response was detected and then decreased to the next lower value. The 50% likelihood of withdrawal was calculated using the up-down von Frey method (Chaplan et al. 1994).

### Rotarod test

The rotarod apparatus (Panlab Harvard Apparatus, Cornella, Spain) was set up in acceleration mode with a range of 4 to 40 rpm such that the maximum speed was reached in 300 seconds. The mice were trained 3 times a day for 5 minutes each for 3 days to learn the task. On test day, the mice were gently placed on the rotarod, and the rotarod was started at the initial speed of 4 rpm. The speed was increased until the mice fell from the moving rotarod, at which point the time was recorded. The trial was repeated 3 times for each mouse, and the mean time was recorded.

### Tissue collection

After the nociceptive and rotarod testing, the mice were deeply anesthetized using 3-4% isoflurane and humanely killed. The L3 and L4 DRG tissues were immediately collected after euthanasia by using the hydraulic extrusion method described earlier (Richner et al. 2017).

### RNA-Sequencing

Total RNAs were extracted from the L3 and L4 DRG tissues using TRIzol (Thermo Fisher Scientific, #15596026). The RNA concentration was quantified using a NanoDrop 1000 (Thermo Fisher Scientific). Good quality of RNA was confirmed using 260nm/280nm absorbance of 2 using a Nanodrop spectrophotometer and was then sent to the Advanced Technology Genomics Core at MD Anderson for RNA-sequencing. We trimmed the RNA sequencing data for low-quality reads using trim_galore, then mapped using STAR onto the mouse genome build mm10. Gene expression was determined using feature Counts (Liao, Smyth, and Shi 2014). Differentially expressed genes were determined after normalization with Remove Unwanted Variation (Risso et al. 2014) using the EdgeR R package.

Enriched pathways were inferred using Gene Set Enrichment Analysis (GSEA) (Subramanian et al., 2005) against pathway compendia REACTOME and Gene Ontology Biological Processes as compiled by the Molecular Signatures Database (MSigDB) (Liberzon et al. 2015). Pathways were considered significant at a false discovery rate < 0.25 per the GSEA developer’s best practices.

### Quantitative RT-PCR

One microgram of RNA was first treated with RNAase-free DNase and then used for reverse transcription with the SuperScript IV VILO Master Mix reverse transcription kit (11766050; Thermo Fisher Scientific). Two microliters of complementary DNA was diluted 5 times and added to a 20-µL reaction volume with SYBR Green real-time PCR mixture (Thermo Fisher Scientific, #A25778). Real-time PCR was performed using the QuantStudio 6 Flex real-time PCR system (Applied Biosystems, Waltham, MA). The thermal cycling conditions were as follows per the manufacturer’s instructions: 1 cycle at 50 °C for 2 minutes, 1 cycle at 95 °C for 2 minutes, and 40 cycles at 95 °C for 15 seconds and at 60 °C for 45 seconds. The primers used are shown in Table 1.

**Table 1.**
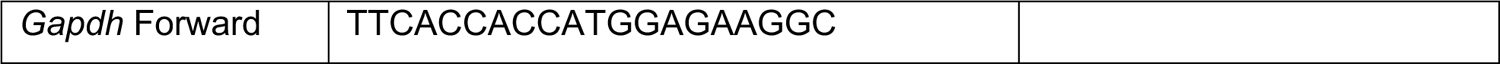
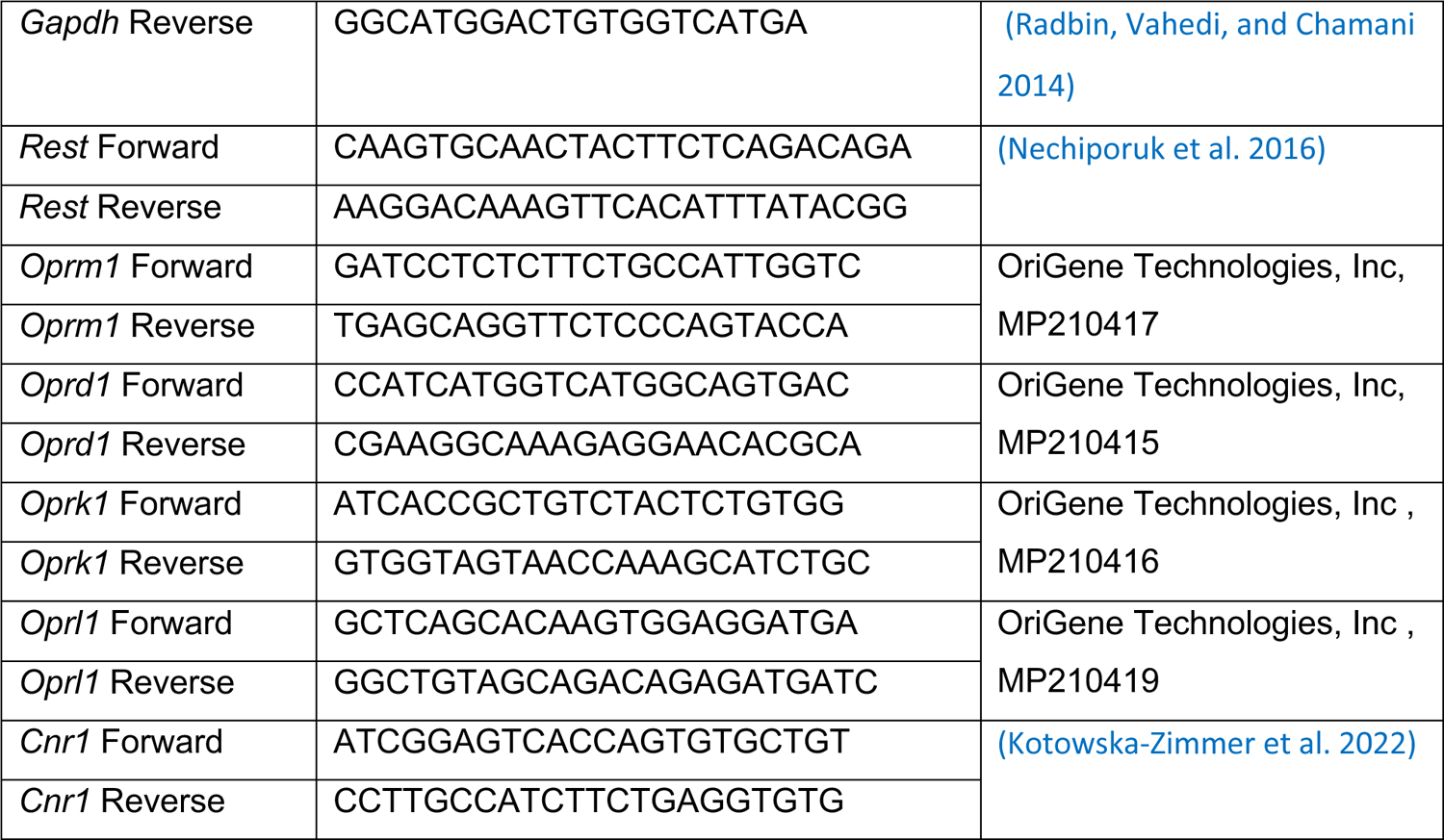
List of primers used in qRT-PCR assays.

### Western Blotting

Protein lysate was made from L3/L4 DRGs using RIPA (89900, Thermo Scientific) added with protease and phosphatase inhibitor (78440, Thermo Scientific). Protein concentration was determined by BCA kit (23228, Thermo Scientific). 20 ugm of protein was loaded and separated on NuPAGE 4 to 12%, Bis-Tris gel (NP0335BOX, Thermo Scientific) under reducing condition in MES SDS running buffer (NP002, Thermo Scientific) as per manufacturer recommendation. Running time was 45 mins at 200V. Transfer was done on nitrocellulose membrane (1620112, Bio Rad) for 90 mins at 120V. The membrane was blocked by 5% skim milk solution prepared by 0.05% Tween 20 in Tris Buffer Saline (TBST) for 1 hour, The primary antibodies were probed for overnight at 4 °C. Membrane washed with TBST for 3 times each and incubated with IRDye 800CW (926-32213, 1:15000, Licor) and IRDye 680RD (926-68072, 1:15000,Licor) for 1 hour. After 3 washes with TBST, membrane was observed using ChemiDOC MP imaging system (Biorad). Rabbit Anti-REST (07579, 1:250, Millipore) and mouse Anti GAPDH (MAB374, 1:300, Millipore) control antibodies were used to probe the membrane.

### ChIP assays

We used the Match tool in the geneXplain platform (https://genexplain.com/genexplain-platform/), which uses the TRANSFAC 2.0 database (version 2023.2) to predict potential REST binding sites in the genome based on the positional weight matrices of binding sites in the database. The promoters of mouse *Oprm1* and *Cnr1* genes were screened from −1000 bp to +1000 bp relative to the transcriptional start site. Next, ChIP-qPCR primers were designed using the NCBI Primer-BLAST tool (https://www.ncbi.nlm.nih.gov/tools/primer-blast/).

We collected L3/L4 DRG tissues from 5 animals (10 DRGs) per group. qChIP was performed using Covaris truChIP (#520237) and Cell Signaling SimpleChIP (#56383) kits as described by the manufacturer with some modifications. Briefly, DRGs from both sham and SNI groups (n=10 from 5 mice, L3 and L4) were crosslinked by 1X fixing buffer A and PFA for 10 minutes. The nuclear fraction was isolated and subjected to sonication (Diagenode Bioruptor PICO B01080010) to break the DNA into 200-700 bp fragments (4 cycles 30/30 on/off time, easy mode). After fragmentation, the chromatin was incubated with either 15 µg REST (Millipore, #07-579 lot#3455213) or 1 µg IgG (Sigma, #27290) antibodies overnight at 4 °C with rotation. The eluted protein-DNA mixture was de-crosslinked using 5 M NaCl and Proteinase K overnight at 65 °C on a Thermomixer. Finally, the DNA was purified using a Qiagen PCR purification kit (#28104) according to the manufacturer’s protocol. The DNA was then subjected to qRT-PCR assay with the primers listed in the following table.

**Table 2.**
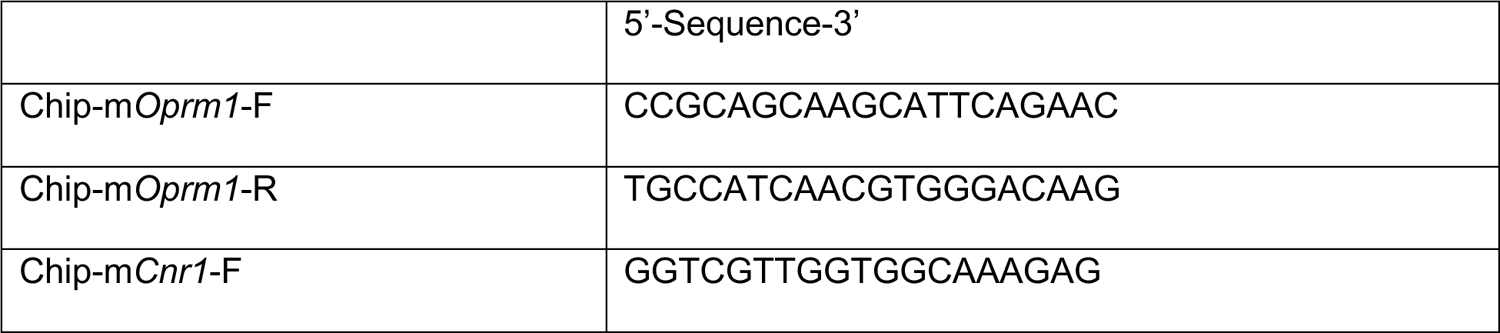
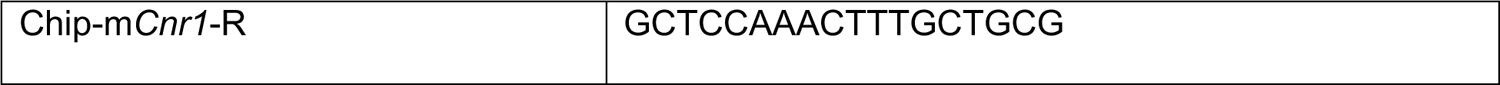
List of primers used in ChIP assays.

Student *t* test was used to compare the groups. Data were expressed as mean +/ S.E.M

## Acknowledgements

We would like to thank Gail Mandel for providing us with the Rest full-length cKO mouse line. This work was partially supported by grants from the National Institutes of Health (R01NS112280 to HLP, SM, and CC; R61DA049334 to HLP and SM). CC was partially supported by CPRIT Core Facility Support Award RP120092.

## Authorship Statement

AS, AT, AEF, YH, BC, SMM, YL, and DG performed experiments, analyzed the data, and wrote corresponding Methods sections, KG and SKS provided methodological help, ER, SG and CC performed bioinformatic data analyses, HLP and SM conceived the project and wrote the manuscript with help from all authors.

## Supplementary Figure Legends

**Supplementary Figure S1.**
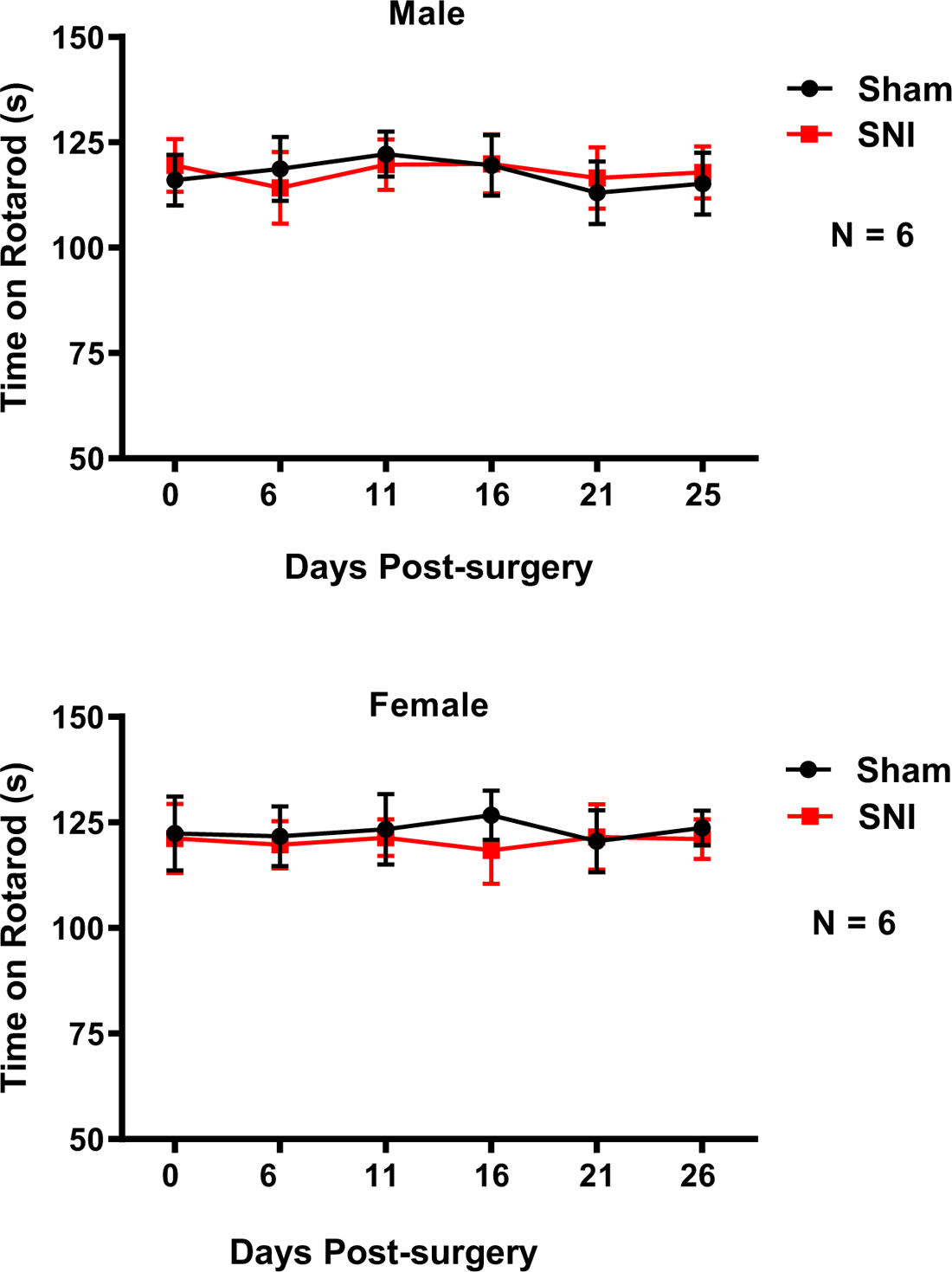
Rotarod assays of wild-type male and female mice indicate no significant difference between sham and SNI treatment. These experiments were performed as described in the Methods section. The trial was repeated 3 times for each mouse, and the mean time was recorded. n= 6.

**Supplementary Figure S2.**
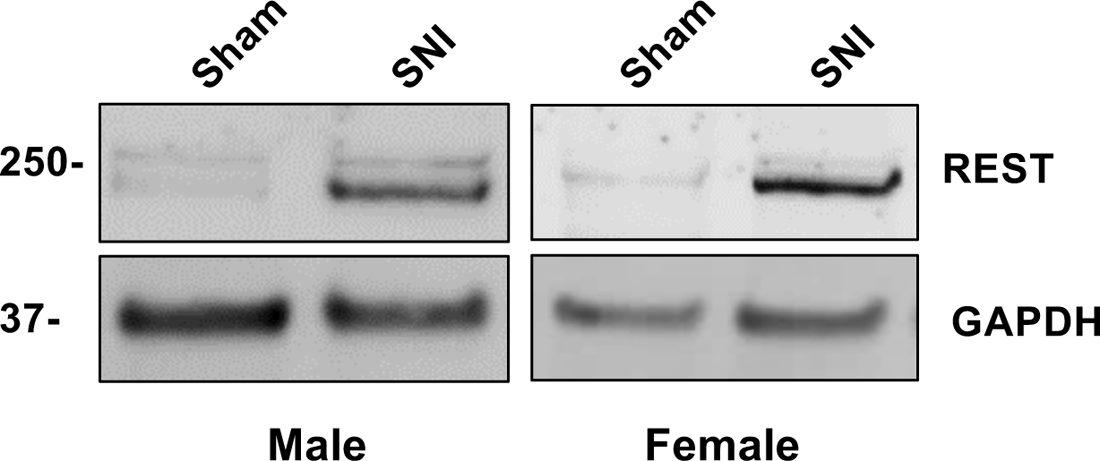
Nerve injury increases REST proteins in the DRG of both male and female wild-type mice. Western blotting was performed from proteins isolated from L3/L4 DRGs of either sham or nerve-injured wild-type male and female mice.

**Supplementary Figure S3.**
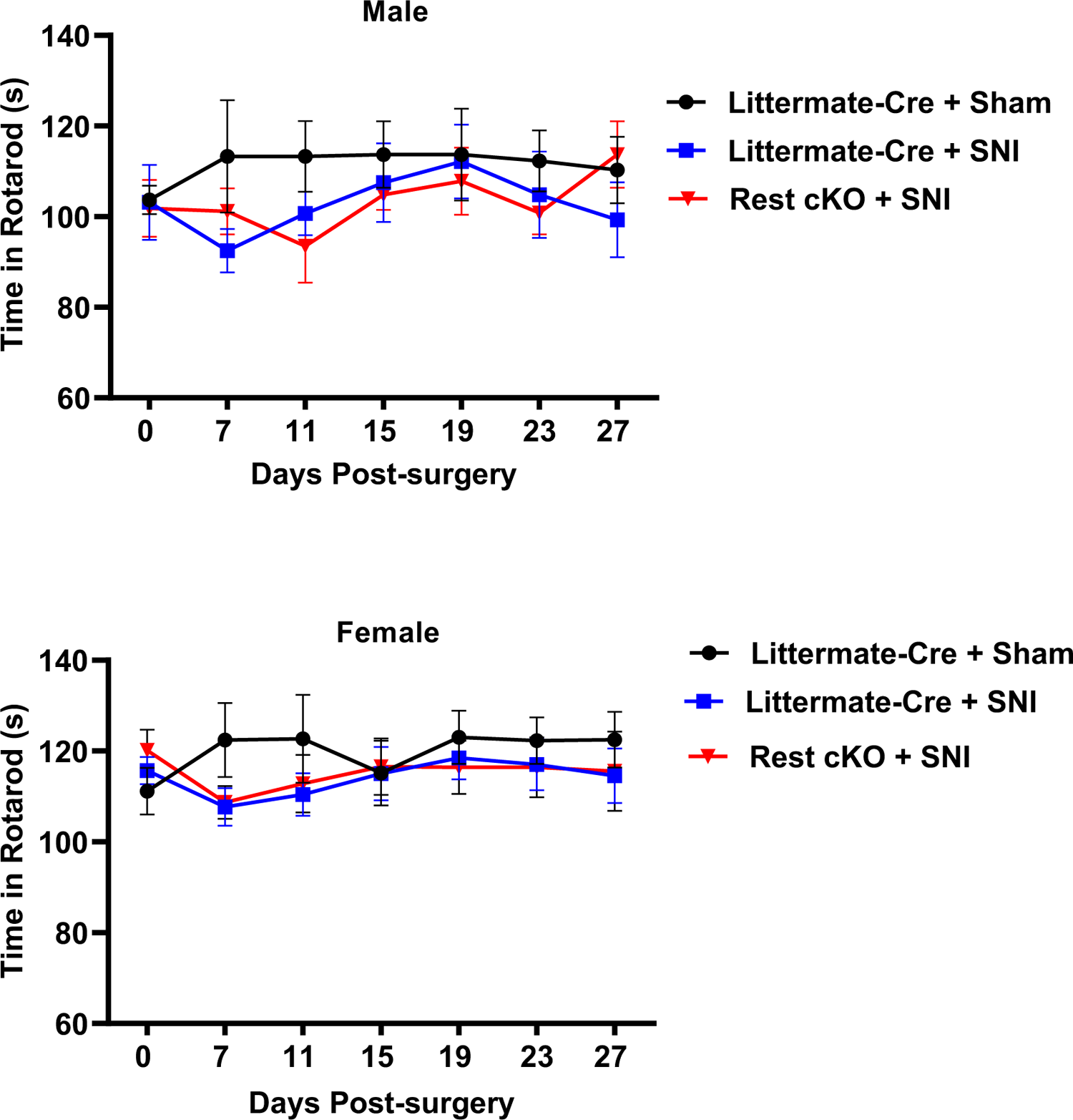
Rotarod assays of littermates (-Cre) + sham, littermates (-Cre) + SNI, and Rest cKO (+Cre) + SNI in male and female mice indicate no significant difference. These experiments were performed as described in the Methods section. n= 6.

**Supplementary Figure S4.**
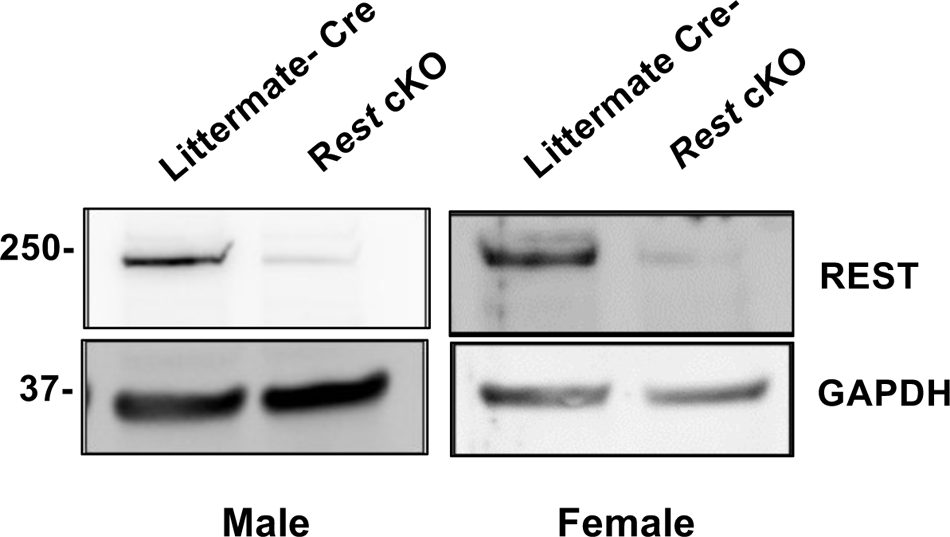
Substantial loss of REST proteins in the DRG of both male and female *Rest* cKO mice. Western blotting was performed from protein isolated from L3/L4 DRGs of *Rest* cKO (+Cre) and littermate (-Cre) male and female mice.

